# Membrane vesicles can contribute to cellulose degradation by *Teredinibacter turnerae,* a cultivable intracellular endosymbiont of shipworms

**DOI:** 10.1101/2024.03.27.587001

**Authors:** Mark T. Gasser, Annie Liu, Marvin Altamia, Bryan R. Brensinger, Sarah L. Brewer, Ron Flatau, Eric R. Hancock, Sarah P. Preheim, Claire Marie Filone, Dan L. Distel

## Abstract

*Teredinibacter turnerae* is a cultivable cellulolytic Gammaproteobacterium (Cellvibrionaceae) that commonly occurs as an intracellular endosymbiont in the gills of wood-eating bivalves of the family Teredinidae (shipworms). The genome of *T. turnerae* encodes a broad range of enzymes that deconstruct cellulose, hemicellulose, and pectin and contribute to wood (lignocellulose) digestion in the shipworm gut. However, the mechanisms by which *T. turnerae* secretes lignocellulolytic enzymes are incompletely understood. Here, we show that *T. turnerae* cultures grown on carboxymethyl cellulose (CMC) produce membrane vesicles (MVs) that include a variety of proteins identified by LC-MS/MS as carbohydrate-active enzymes (CAZymes) with predicted activities against cellulose, hemicellulose, and pectin. Reducing sugar assays and zymography confirm that these MVs exhibit cellulolytic activity, as evidenced by the hydrolysis of CMC. Additionally, these MVs were enriched with *TonB*-dependent receptors, which are essential to carbohydrate and iron acquisition by free-living bacteria. These observations indicate a potential role for MVs in lignocellulose utilization by *T. turnerae* in the free-living state, suggest possible mechanisms for host-symbiont interaction, and may be informative for commercial applications such as enzyme production and lignocellulosic biomass conversion.

## Introduction

The biological degradation of lignocellulose, a primary component of wood and all plant biomass, is a critical process in the global carbon cycle and has potential applications in renewable energy and chemical production through biomass conversion (Ragauskas *et al*., 2014; Østby *et al*., 2020). Lignocellulose is a complex composite material composed of microfibrils of cellulose, the most abundant organic polymer on Earth, embedded in a matrix of hemicellulose, pectin, and lignin (Cragg *et al*., 2015; Dahmen *et al*., 2019). However, due to the structural complexity of lignocellulose, its degradation typically requires multiple enzymes to disassemble and convert its component polymers into accessible nutrients. This complexity limits lignocellulose as a carbon source for most animals and creates obstacles to bioconversion to renewable fuels or fine chemicals (Cragg *et al*., 2015). Nonetheless, many animals, including herbivores, termites, and wood-boring bivalves (shipworms), can digest lignocellulose in leafy or woody plant material with the aid of hydrolytic enzymes produced by symbiotic bacteria (Cragg *et al*., 2015).

Bacteria use a variety of mechanisms to degrade lignocellulosic substrates. For example, polysaccharide-degrading bacteria, including *Bacteroides*, *Clostridia*, and *Fibrobacter,* commonly found in the gut microbiomes of ruminants, use the type IX secretion pathway (T9SS) to export large proteins, including multidomain carbohydrate-active enzymes (CAZymes) (Gharechahi *et al*., 2023). These enzymes may be released into the environment or bound to the cell envelope (McGavin *et al*., 1990; Cai *et al*., 1999; Yan and Wu, 2013). Rumin bacteria may also export CAZymes via membrane vesicles (MVs) (Forsberg *et al*., 1981; La Rosa *et al*., 2022; Gharechahi *et al*., 2023). MVs are membrane-enclosed spheres, typically 40-400 µM in diameter, that may form by blebbing or pinching off the outer membrane of diderm bacteria (Toyofuku *et al*., 2023). MVs formed by this mechanism are called Outer Membrane Vesicles (OMVs) and contain mainly periplasmic and outer membrane components (Guerrero-Mandujano *et al*., 2017). Bacteria may also form MVs by other mechanisms, resulting in MV types that contain cytoplasmic membranes and other cytoplasmic components (Schwechheimer and Kuehn, 2015; Nagakubo *et al*., 2020; Huang *et al*., 2022; Toyofuku *et al*., 2023). Multiple MV types may be produced simultaneously by the same bacterium (Gan *et al*., 2023).

Bacterial MVs have been implicated in a wide variety of functions, including the export of proteins, lipids, carbohydrates, metabolites, and nucleic acids; waste disposal; nutrient scavenging; defense against phage and antibiotics; and delivery of bioactive molecules across the plasma membranes of animal hosts (Toyofuku *et al*., 2023). However, the extent to which MVs contribute to bacterial lignocellulose degradation remains incompletely explored, particularly in environments other than ruminant guts. In the rumen, members of several bacterial phyla, including Fibrobacterota (Arntzen *et al*., 2017), Bacilota (Ichikawa *et al*., 2019), and Bacteroidota (Forsberg *et al*., 1981), produce MVs highly enriched in CAZymes capable of degrading cellulose and other plant biomass. However, to our knowledge, no member of the phylum Pseudomonadota (formerly Proteobacteria), the largest and most phenotypically diverse bacterial phylum (Rosenberg *et al*., 2014), has been shown to produce MV’s with a protein content that suggests specialization for degrading lignocellulose.

One example of an ecologically important lignocellulose-degrading Pseudomondadota species is *Teredinibacter turnerae.* This gammaproteobacterium is a member of the family Celvibrionaceae, which includes many polysaccharide-degrading marine bacteria (Spring *et al*., 2015). *T. turnerae* is unusual in that it can exist either as a versatile free-living marine lignocellulose-degrading bacterium (Distel *et al*., 2002) or as an intracellular symbiont of shipworms (Distel *et al*., 1991), a family of bivalve mollusks that are the primary consumers of wood in marine environments (Voight, 2015; Cragg *et al*., 2020). Interestingly, unlike other marine Cellvibrionaceae, the *T. turnerae* genome includes a broad representation of genes associated with the degradation of terrestrial woody plant materials, including cellulose, xylan, mannan, galactorhamnan and pectin, but lacks enzyme systems for degradation of common marine polysaccharides including agar, alginate, and fucoidan and has only comparatively sparse representation of chitinase and laminarinase genes (Yang *et al*., 2009).

As a free-living bacterium, *T. turnerae* grows on both soluble and insoluble cellulosic substrates, including carboxymethylcellulose (CMC), short-chain cellooligosaccharides, cellobiose, purified cellulose, e.g., Sigmacell or Avicel cellulose), paper, e.g., Whatman #1 filter paper (Waterbury *et al*., 1983; Distel *et al*., 2002), and crude plant biomass, e.g., pine sawdust, cotton fiber, coconut hull fiber, and rice hulls (unpublished observation).

As an intracellular symbiont, *T. turnerae* grows within specialized cells called bacteriocytes found in the gland of Deshayes, a tissue located within the shipworm’s gills (Popham and Dickson, 1973; Popham, 1974; Distel *et al*., 1991). The genome of *T. turnerae* and other shipworm symbionts from the genus *Teredinibacter* encode a wide variety of enzymes targeting lignocellulose (Distel *et al*., 2002; Yang *et al*., 2009; O’Connor *et al*., 2014; Brito *et al*., 2018; Altamia *et al*., 2020, 2021) and express them within this gland (O’Connor *et al*., 2014; Sabbadin *et al*., 2018). Enzymes encoded in the shipworm symbiont’s genomes are also found in the shipworm’s cecum (O’Connor *et al*., 2014; Sabbadin *et al*., 2018; Altamia and Distel, 2022). This gut compartment is a primary location of wood digestion in shipworms and is nearly devoid of microbes (Betcher *et al*., 2012). The distribution of symbiont-made enzymes in both the intracellular environment of the host and in the gut, which is part of the host’s external environment, indicates the existence of an unusual and unexplained protein transport system that potentially exports symbiont-made proteins across the host’s cellular membranes (Altamia and Distel, 2022).

While explaining the complex interaction of *T. turnerae* with its shipworm host is beyond the scope of this investigation, understanding the mechanisms by which this bacterium exports lignocellulolytic enzymes in the free-living state may shed light on potential mechanisms by which these enzymes may be transported within its shipworm hosts. MVs are a versatile system through which bacteria exchange proteins and other biomolecules with their external environment and the internal environment of their hosts (Gan *et al*., 2023; Toyofuku *et al*., 2023). Therefore, we ask whether free-living cells of *T. turnerae* export lignocellulose-degrading enzymes via MVs.

To address this question, we purify and characterize the protein composition of MVs produced by *T. turnerae* during growth on carboxymethyl cellulose (CMC), a water-soluble derivative of cellulose. We identified proteins in the purified MVs that are putatively involved in interactions with lignocellulose by using LC-MS/MS analysis and comparison to the Carbohydrate Active Enzyme Database (CAZy) (Lombard *et al*., 2014). Additionally, we performed activity assays to show that MVs produced by *T. turnerae* are capable of cellulose hydrolysis. These observations suggest that MVs contribute to lignocellulose utilization by *T. turnerae* in the free-living state and provide clues that might help explain enzyme transport during host-symbiont interaction.

## Experimental Procedures

### Strain isolation, selection, and growth conditions

Two strains of *Teredinibacter turnerae* were selected for this study: *strain* T7901 (ATCC 39867; GenBank CP001614) and a closely related strain SR01903 (GenBank CP149818). The former was isolated from the shipworm *Bankia gouldi* collected in Beaufort, NC, in 1979 and was the first shipworm symbiont strain brought into pure culture (Waterbury *et al*., 1983; Distel *et al*., 2002; Yang *et al*., 2009). The latter was isolated from the gills of a single specimen of *Lyrodus pedicellatus* found in naturally occurring driftwood collected by hand in shallow water in the Indian River Lagoon, Merit Island, FL. (N 28.40605 W 80.66034) on January 24, 2020. Gills were removed by dissection and homogenized in 1.0 ml of shipworm basal medium (SBM) (Waterbury *et al*., 1983) in an autoclave-sterilized glass dounce homogenizer. Homogenate was streaked onto a culture plate containing 1.0% Bacto agar and SBM at pH 8.0 supplemented with 0.2% w/v powdered cellulose (Sigmacell Type 101; Sigma-Aldrich) and 0.025% NH_4_Cl. Plates were incubated at 30 °C until individual colonies could be observed. An individual colony was then picked and subjected to multiple rounds of restreaking from single colonies to ensure clonality. We PCR-amplified and sequenced the 16S rRNA gene to identify this isolate and compared it to published *T. turnerae* genomes. The complete genome of *T. turnerae* SR01903 was then sequenced (SRA number SRR28421271) and submitted to Genbank (CP149818). For the experiments described herein, strains were initially propagated in 6 mL cultures in SBM supplemented with 0.025% NH_4_Cl and 0.2% carboxymethyl cellulose for 4 days before being diluted 1/250 in fresh media and harvested after 2 days (OD600 0.2-0.3). All cultures were incubated in a shaker incubator at 30 °C and 100 rpm. An overview of the procedures conducted in this research is presented in Figure 1.

**Figure 1.**
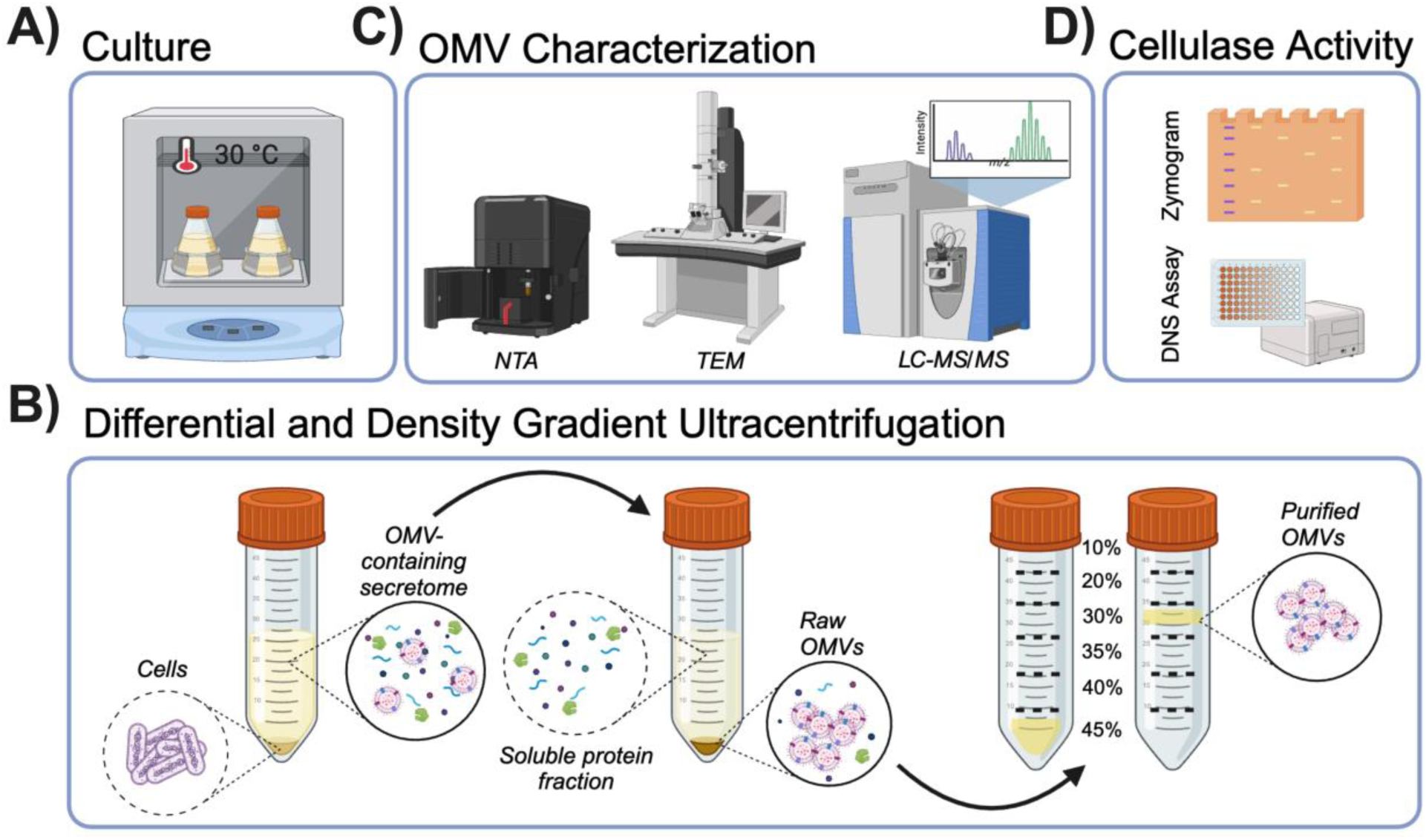
Methods used to isolate and characterize membrane vesicles (MVs) from *T. turnerae*. Diagram showing: (A) bacterial culture, (B) MV isolation based on differential and density gradient separation, (C) MV visualization and size determination by TEM and NTA, proteome analysis by LC-MS/MS, and (D) detection of cellulase activity in purified MV preparations by zymography and reducing sugar (DNS) assay. Created with BioRender.com

### Isolation of Membrane Vesicles (MVs)

Bacterial cells were separated from the culture supernatant by centrifugation at 5,000 x g for 20 minutes at 4 °C. The supernatant was then transferred to a fresh tube, and centrifugation was repeated to remove residual bacterial cells. The final supernatant was carefully collected without disturbing the remaining pellet and filtered through a 0.22 μm polyethersulfone filter as an additional precaution to remove cells. The putative MVs were then pelleted from the filtrate by ultracentrifugation at 120,000 x g (T-647.5 rotor, Sorvall) for 90 minutes at 4 °C. For purification, the resulting pellet was resuspended in 0.1 M phosphate-buffered saline (PBS) and fractionated by bottom-up density gradient ultracentrifugation using Optiprep™ (iodixanol density gradient medium) as follows. The resuspended MV-containing pellet was mixed with 60% iodixanol solution to a final density of 45% (w/v) and placed at the bottom of an ultracentrifuge tube. A discontinuous density gradient was generated using a syringe and G21 needle to deposit 40%, 35%, 30%, 20%, and 10% iodixanol layers with a final layer of 0.25 mL 0.1 M PBS on top. The resulting gradient was centrifuged for 16 hours at 150,000 x g and 4 °C (SW55 Ti rotor, Beckman-Coulter). Sample banding was visually observed at the interface of the 30% and 20% fractions (Supporting Information: Figure S1). All fractions were collected, and iodixanol was removed by passive diffusion dialysis (1,000 kDa MWCO) in exchanging buffer of 50 mM ammonium bicarbonate pH 8.3 at 4 °C. MV samples were removed from the dialysis bag and concentrated under vacuum to 100 μL (SPD121P SpeedVac, Thermo Scientific). Particle visualization and characterization proceeded with the single 30% fraction.

### Transmission electron microscopy (TEM)

Purified MVs were diluted in molecular grade distilled water and absorbed onto 200 mesh formvar-treated and carbon-coated copper grids for 60 seconds. Samples were then fixed for 5 minutes in 4% glutaraldehyde in 0.1 M sodium cacodylate, and grids were negatively stained with 1% aqueous uranyl acetate for 60 seconds and left to dry. MVs were imaged in an FEI Tecnai T12 (Thermo Fisher) transmission electron microscope at 80 kV with an AMT bottom-mount camera.

### Nanoparticle tracking analysis (NTA)

NTA was performed using the ZetaView® PMX-220 Twin (Particle Metrix) configured with 488 nm and 640 nm lasers with long wave-pass cut-off filters (500 nm and 660 nm, respectively) and a sensitive CMOS camera 640 x 480 pixels. Samples were diluted in 2 mL of 0.1 μm filtered deionized water (18 MΩ/cm) to obtain a particle concentration between 1 x 10^7^ and 1 x 10^8^ particles/mL. The instrument was set to a sensitivity of 80, a shutter speed of 100, and a frame rate of 30 frames per second. Each sample was measured at 11 different positions throughout the sample cell, with 1 cycle of reading at each position to have a minimum of 1,000 traces. If the number of traces was below 1,000 counts, some additional sample was flushed inside the sample cell, and the acquisition was repeated. Post-acquisition parameters were set to a minimum brightness of 20, a maximum size area of 1,000 pixels, a minimum size area of 10 pixels, and tracelength of 15 frames. Automated cell quality control was checked using high-quality deionized water. Camera alignment and focus optimization were performed using polystyrene Nanosphere™ 100 nm size standard beads. Data analysis was performed with ZetaView® 8.05.14 software provided by the manufacturer. Automated reports of the particle recordings across the 11 positions were manually checked, and any outlier position was removed to calculate particle concentration and distribution.

### Proteomic analysis

MVs were lysed and denatured in a solution containing 5% sodium dodecyl sulfate (SDS), 100 mM Tris (pH=8), 20 mM chloroacetamide, and 10 mM tris (2-carboxyethyl) phosphine hydrochloride and incubated at 90 °C for 10 minutes. Proteins were aggregated and isolated using Sera-Mag™ carboxylate-modified SpeedBeads (SP3 beads) and digested in 100 mM NH_4_HCO_3_ containing Trypsin (final concentration 6 ng/μL) overnight at 37 °C and 1,200 rpm. Digested samples were loaded onto C18 tips, and peptides were separated online using an 88-minute LC gradient (Evosep LC system). MS analysis was performed on a Q-Exactive HF-X Hybrid Orbitrap mass spectrometer (Thermo Scientific). One full high-resolution MS spectrum was acquired with a resolution of 45,000, an AGC target of 3 x 10^6^ with a maximum ion time of 45 ms, and a scan range of 400-1,500 m/z, followed by 20 HCD MS/MS scans with a resolution of 1,5000, an AGC target of 1 x 10^5^ with a maximum ion time of 120 ms, NCE of 27, and isolation window of 2 m/z. The resulting MS/MS spectra were searched against the strain-specific and universal contaminant protein databases using Sequest within Proteome Discoverer 1.4. Identifications were filtered to include only high-confidence peptides based on a less than 1% false discovery rate (FDR) against a decoy database. FDR estimates the expected ratio of false positive classifications (false discoveries) to total positive classifications. Proteins were linked to peptides, and only proteins with >2 peptide spectral matches (PSM) with two unique peptides were kept (Supporting Information: Dataset S1).

### Functional characterization of MV proteins

Identified proteins were annotated against the Clusters of Orthologous Groups (COG) of proteins database (Galperin *et al*., 2021). Additionally, protein subcellular locations and secretion signals were predicted using DeepLocPro v1.0 (https://services.healthtech.dtu.dk/services/DeepLocPro-1.0/) (Moreno *et al*., 2024) and Signal v5.0 (https://services.healthtech.dtu.dk/services/SignalP-5.0/) (Almagro Armenteros *et al*., 2019), respectively. Enrichment and functional associations between proteins were determined by STRINGdb v11 (https://string-db.org/) (Szklarczyk *et al*., 2019) and ShinyGO v0.8 (http://bioinformatics.sdstate.edu/go/) (Ge *et al*., 2018). Settings were selected to use all coding sequences in the T7901 genome as background and a minimum FDR stringency of 1 x 10^-5^. We then established which MV proteins are predicted to exhibit carbohydrate-active properties using the annotated T7901 genome in the Carbohydrate Active Enzymes Database (CAZy, https://cazy.org) (Lombard *et al*., 2014).

### Enzymatic activity assays

The cellulolytic activity of proteins in MV preparations was visualized and size fractionated by denaturing polyacrylamide gel electrophoresis (SDS-PAGE, 1.5 mm thickness, 9% acrylamide, and 0.2% CMC final concentration). For zymographic analysis, 14 µg of protein sample determined by Pierce 660 assay (Thermo Scientific) using bovine serum albumin as standard were heat denatured by boiling for 3 minutes in SDS without a reducing agent to facilitate recovery of activity after refolding and added to each lane. After electrophoresis, the SDS-PAGE gel was transferred to 500 mL of refolding buffer (20% isopropanol, 0.1% Triton X-100 in 1x PBS, pH 7.4) and gently shaken for 1 hour at room temperature. The gel was then incubated in fresh 1x PBS buffer, pH 7.4, for 16 hours at room temperature before being stained with 0.1% Congo Red in 1x PBS buffer, pH 7.4 for 1 hour. The zymogram was destained with 1 M NaCl overnight to visualize regions where cellulase activity removed CMC.

Additionally, the production of reducing ends was measured by the colorimetric 3,5-dinitrosalicylic acid (DNS) method (Ghose, 1987). Reactions were carried out in 96-well plates containing 50 μL 1% CMC, 30 μL 50 mM citrate buffer, and 20 μL purified MV suspension and incubated for 4 hours at 37 °C. Next, 900 μL of DNS was added to each reaction, and reactions were heated at 99 °C for 10 minutes. Absorbance was measured at 540 nm, and sugar concentration was calculated by comparison to a glucose standard curve. The enzyme activity unit was defined as the amount of enzyme that liberated 1 μmol of reducing sugar per minute.

## Results

### MV isolation and characterization

MVs were isolated from cell-free supernatants of two closely related strains of *T. turnerae*, T7901 and SR01903, by density gradient ultracentrifugation. The presence of MVs was confirmed by NTA and TEM (Figure 2 A-D). The mean concentration of particles and average particle diameter determined by NTA were 1.67 x 10^9^ particles/mL and 162.5 ± 49.7 nm for T7901 and 1.30 x 10^9^ particles/mL and 206.6 ± 61.7 nm for SR01903. TEM images (Figure 2 C-D) show that the putative MV-enriched fractions from strains T7901 and SR01903 contain particles with size distribution and morphology typical of bacterial MVs.

**Figure 2.**
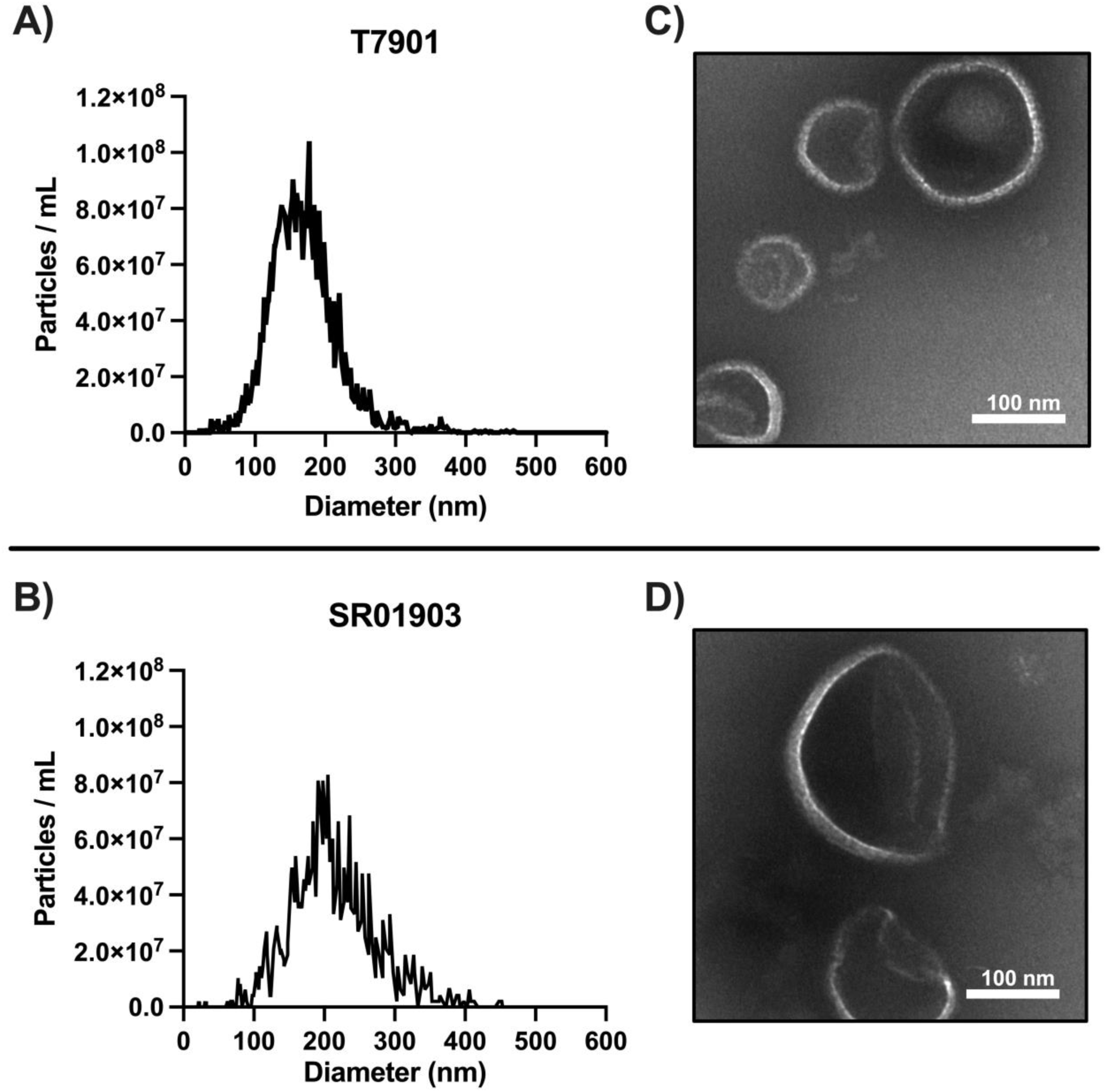
Size, concentration, and visualization of MVs isolated from *T. turnerae* culture supernatants after growth on carboxymethyl cellulose (CMC). (A-B) Size distribution and particle concentration for strains T7901 and SR01903, respectively. MVs from SR01903 were slightly larger in average size than MVs from T7901. (C-D) Isolated MVs from strains T7901 and SR01903, respectively, were imaged using transmission electron microscopy. Scale bar = 100 nm.

### MV proteome analysis

The protein content of MVs from *T. turnerae* strains T7901 and SR01903 were analyzed by LC-MS/MS. Peptide-generated data were used to search against T7901 and SR01903 annotated protein sequences to identify MV proteins and their relative abundance. A cut-off of two or more unique predicted peptides per identified protein was used, identifying 472 and 569 proteins for T7901 and SR01903, respectively. The Top 100 most abundant proteins identified in T7901 and SR01903 represented 81% and 74% of the total peptide content, respectively. Among those proteins, 71 were identified in both strains, indicating that the two strains produce a shared core set of MV proteins when grown under the examined culture condition. After protein identification, DeepLocPro v1.0 (Moreno *et al*., 2024) and Signal v5.0 (Almagro Armenteros *et al*., 2019) were used to predict the subcellular location and secretion signals of all identified proteins.

Outer membrane proteins comprised most (∼76% and ∼71%, respectively) of the MV protein mass of both T7901 and SR01903 (Figure 3 A). Although cytoplasmic and inner membrane proteins were detected, these constituted only about 9% of total protein by mass. This indicates that these MV preparations were mainly comprised of outer membrane vesicles but may have contained other types, e.g., outer inner membrane vesicles (OIMVs) (Toyofuku *et al*., 2023).

**Figure 3.**
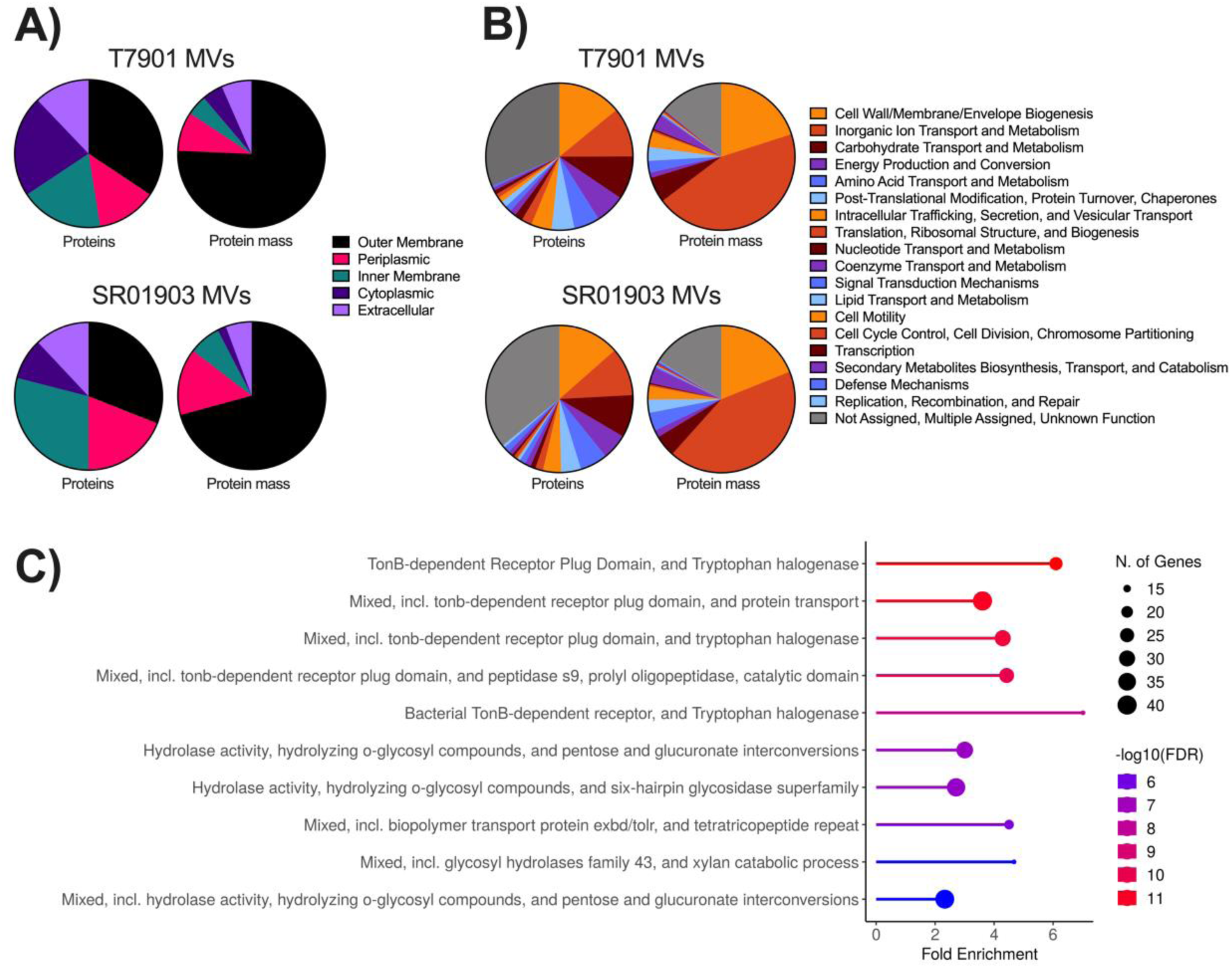
T. turnerae MV proteome profiles. (A) Relative protein abundance grouped by predicted subcellular location and (B) Clusters of Orthologous Groups (COG) functional categories of identified proteins. Six color-safe hues represent COG categories and are repeated in order starting from 12 o’clock on the graph and moving clockwise. (C) Top 10 STRINGdb pathways ranked by false discovery rate (FDR) found in MVs of T7901. Similar top pathways were found in the MVs of SR01903. STRING analysis was performed against the corresponding bacterial strain genome background to estimate the amount or fold of enrichment, i.e., the representation of that functional category above what is expected by chance. FDR estimates the expected ratio of false positive classifications (false discoveries) to total positive classifications.

MV proteins were clustered into orthologous groups (COG). Most MV proteins assigned to clusters in both strains were categorized as Cell Wall/Membrane/Envelope Biogenesis, Inorganic Ion Transport and Metabolism, and Carbohydrate Transport and Metabolism (Figure 3 B).

### *TonB*-dependent receptors and glycosyl hydrolase activity are enriched in MVs

MV protein content was analyzed using STRING (Szklarczyk *et al*., 2019) enrichment to highlight functional associations. We found significant enrichment of pathways involving *TonB*-dependent receptors (TBDR) and glycosyl hydrolase activity (Figure 3 C). The most abundant proteins were TBDRs, which constituted ∼47% and ∼44% of the MV protein content by mass in T7901 and SR01903, respectively. In total, 39 and 48 predicted CAZymes were present, representing 30 and 29 different catalytic CAZy families and 8.3% and 8.4% of identified MV proteins in T7901 and SR01903, respectively. Most identified CAZy modules belong to glycoside hydrolase families (GH) predicted to be involved in cellulose and hemicellulose degradation (Figure 4 A) and that include several hydrolytic activities: for example, endo-glucanases (GH5, GH8, GH9, GH10, GH11, GH26, GH44, GH45, GH51, GH62), exo-glucanases (GH5, GH9, GH26, GH43), and beta-glucosidases (GH2, GH3, GH5, GH39). In addition, multiple enzymes were predicted to include non-catalytic carbohydrate-binding modules (CBMs), with predicted specificity for cellulose or hemicellulose components.

**Figure 4.**
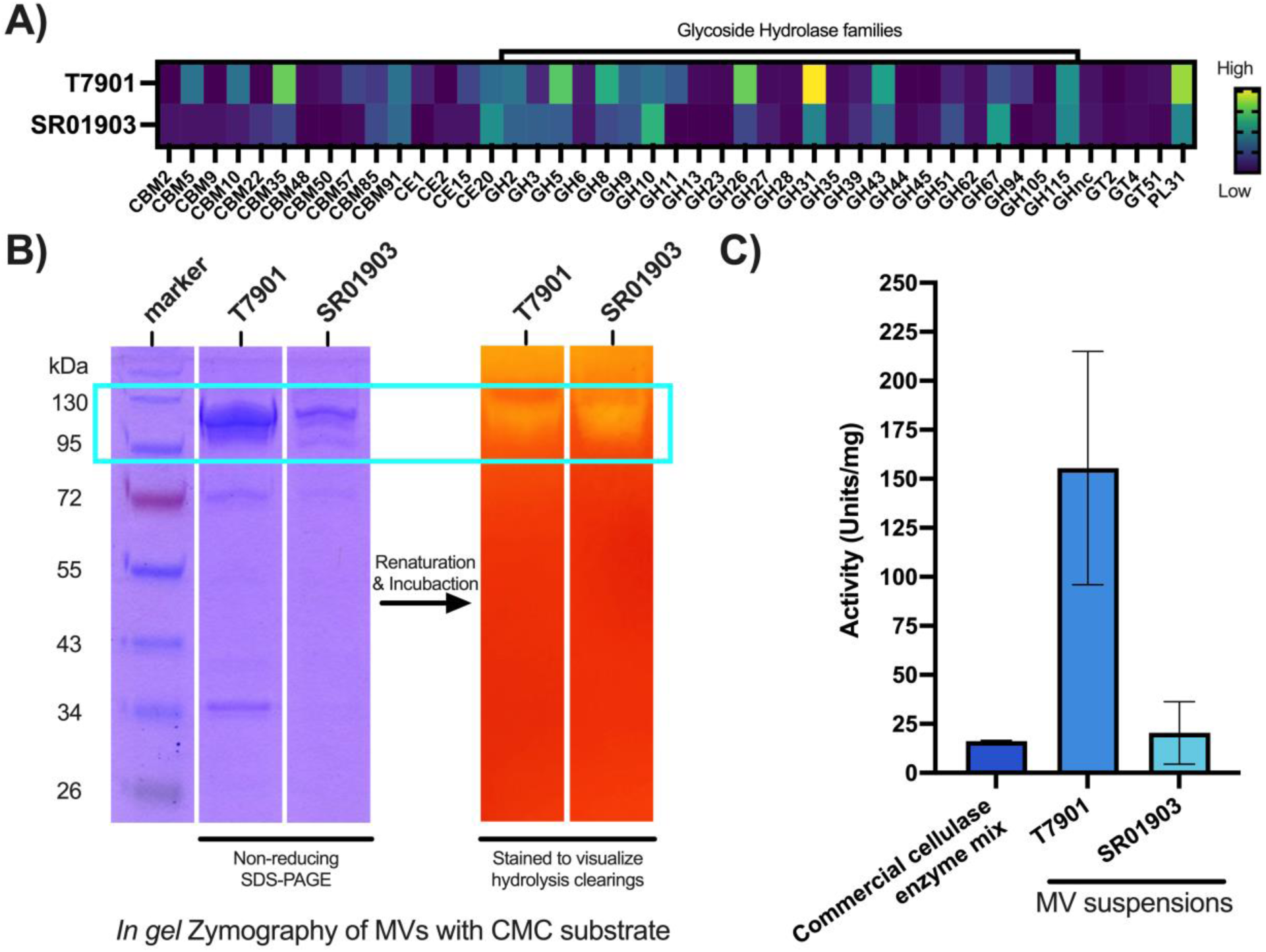
Occurrence and relative abundance of carbohydrate-active modules and detection of cellulase activity in *T. turnerae* MVs. (A) Heatmap profile of the relative abundance of the predicted carbohydrate-active enzymes found in MVs. (B) Denaturing nonreducing SDS-PAGE (left) and zymogram (right; CMC substrate) of isolated MVs. (C) Histogram showing cellulase activity (CMC substrate) in Units/mg (as determined by DNS assay) for isolated MVs and a purified cellulase enzyme cocktail (Cellulase R-10, Goldbio CAS# 9012-54-8). Bars indicate means (error bars: standard deviations of three replicates).

### Cellulase activity

Many MV proteins were predicted to have or be associated with cellulolytic activity. We confirmed these by testing MV suspensions for hydrolytic activity against CMC by zymography and reducing sugar DNS assay. In zymograms, clear zones indicating endoglucanase activity (Teather and Wood, 1982) were visualized between 95-130 kDa for both strains after 16-hour incubation (Figure 4 B). The production of reducing sugars from CMC was measured using DNS assay, and when normalized to total protein content, MVs produced from SR01903 displayed similar or significantly greater cellulolytic activity compared to a commercial soluble cellulase enzyme cocktail (Figure 4C). Notably, MVs produced from T7901 showed ∼10x the activity of the commercial cellulase cocktail (Figure 4 C) against CMC. While we observed a substantial difference in CMC hydrolysis between MVs from the two *T. turnerae* strains, the ecological and physiological significance of this difference is difficult to interpret as CMC is not a naturally occurring substrate, and we did not control for differences in the density beyond OD600 measurement or physiological state of cells at the time of MV purification.

## Discussion

### MVs in bacterial lignocellulose degradation

The extent to which bacterial MVs contribute to lignocellulose degradation and the types of bacteria that produce lignocellulolytic MVs have been investigated in gut microbiomes (Forsberg *et al*., 1981; McGavin *et al*., 1990; Elhenawy *et al*., 2014; Burnet *et al*., 2015; Arntzen *et al*., 2017; La Rosa *et al*., 2022; Gharechahi *et al*., 2023) but are less well explored in other organisms and environments. In bacteria, efficient deconstruction of lignocellulose typically requires multiple carbohydrate-active enzymes (CAZymes), each of which may contain multiple catalytic and carbohydrate-binding modules. These catalytic modules encompass a broad range of enzymatic activities. Glycoside hydrolase (GH) modules may have endo- or exo-activities targeting the inner or terminal bonds of cellulose and hemicellulose backbones, as well as activities targeting the oligomers and monomers that constitute cellulose and hemicellulose. Carbohydrate esterase (CE) modules cleave bonds that form hemicellulose side chains and bonds between hemicellulose and lignin. Carbohydrate binding modules (CBMs) adhere to various carbohydrate components of lignocellulose and can enhance enzyme activity by increasing accessibility to the substrate (Hervé *et al*., 2010).

Packaging of lignocellulose-degrading CAZymes within MVs or bound to MV surfaces can have distinct advantages over their secretion as soluble proteins or proteins bound to the cell surface. For example, MVs may protect proteins from degradation (Bonnington and Kuehn, 2014; Alves *et al*., 2016; Zingl *et al*., 2021), selectively concentrate proteins with specific functions (Orench-Rivera and Kuehn, 2016; Sartorio *et al*., 2023), and deliver proteins to remote locations in sufficient quantity to produce a desired effect without the dilution that soluble proteins would experience (Toyofuku *et al*., 2023). These factors may be significant for the degradation of complex and insoluble substrates like lignocellulose, where the simultaneous delivery of specific sets of catalytic and carbohydrate-binding modules in the correct proportions and spatial orientation can yield significant synergistic interactions and enhance efficiency (Park *et al*., 2014).

Despite these advantages and the ubiquitous production of MVs by bacteria, members of only a few bacterial phyla, notably Fibrobacterota (Arntzen *et al*., 2017), Bacilota (Ichikawa *et al*., 2019), and Bacteroidota (Forsberg *et al*., 1981), have been shown to produce MVs capable of deconstructing lignocellulose. Here, we report that *Teredinibacter turnerae* (Pseudomonadota; Gammaproteobacteria; Cellvibrionacea) produces MVs containing a wide variety of enzymes with predicted activity against lignocellulose and that cellulolytic activity is retained by MVs upon purification.

Among Pseudomonadota, several species package carbohydrate-active enzymes in MVs, many of which target marine plant polysaccharides. For example, when grown on k-carrageenan, the marine gammaproteobacterium *Alteromonas macleodii* produces MVs enriched in CAZymes (Naval and Chandra, 2019). These include GH16, a family whose members are active against multiple marine plant polysaccharides, including agar, k-carrageenan, and porphyrin, GH28 which includes mainly polygalacturonases, and GH42, which are mostly B-galactosidases (Naval and Chandra, 2019). Similarly, *Pseudoalteromonas distincta* produces MVs with predicted hydrolytic activity against pectin and alginate (Dürwald *et al*., 2021).

MVs may also contain enzymes active against components of lignocellulose. For example, the plant pathogen *Xanthomonas campestris* produces MVs containing two virulence-associated xylanases involved in pathogenic infection (Solé *et al*., 2015). In *Pseudomonas putida*, MVs play a role in the degradation of lignin-derived aromatic monomers but not oligomeric lignin (Choi *et al*., 2014; Salvachúa *et al*., 2020).

### Lignocellulolytic MVs produced by *T. turnerae*

When grown in pure culture and within the gills of its shipworm hosts, *T. turnerae* secretes a broad array of CAZymes that specifically target lignocellulose (Yang *et al*., 2009; O’Connor *et al*., 2014; Sabbadin *et al*., 2018). However, the role of MVs in transporting cellulolytic enzymes by *T. turnerae* has not been investigated. Here, we show that lignocellulolytic CAZymes constituted a substantial proportion of the MV proteomes of *T. turnerae* strains T7901 and SR01903 (8% and 6% by count and 8% and 5% by mass, respectively). Of these, 94% have predicted secretion signals, and 43% are predicted to be lipoproteins anchored to the outer membrane by a lipid moiety. Detected MV proteins included representatives of 11 carbohydrate-binding module (CBM) families (CBM2, 5, 9, 10, 22, 35, 48, 50, 57, 85, and 91). Each of these families is known to bind cellulose or hemicellulose components, except for CBM50, which typically binds peptidoglycan or chitin. Additionally, catalytic modules representing 23 glycoside hydrolase (GH) families, four carbohydrate esterase (CE) families, and one polysaccharide lyase (PL) were detected. All of these GH families (GH2, 3, 5, 8, 9, 10, 11, 13, 23, 26, 28, 31, 35, 39, 43, 44, 45, 51, 62, 67, 94, 105, and 115) target bonds found within cellulose, hemicellulose or pectin, except GH23, which primarily acts on peptidoglycan. Finally, the four carbohydrate esterase (CE) families detected (CE1, 2, 15, and 20) target bonds found in hemicellulose, or bonds covalently linking hemicellulose to lignin. Thus, bioinformatic predictions indicate the ability of *T. turnerae* MVs to hydrolyze a wide range of bonds in lignocellulose, including potential endo- and exo-activities against cellulose and hemicellulose backbones, debranching of hemicellulose sidechains, and hydrolysis of the resulting oligomers and monomers.

The fact that this wide array of activities was found in MVs produced by cells of *T. turnerae* strains T7901 and SR01903 grown with pure CMC as a sole carbon source indicates that the expression of genes active against other lignocellulose components was either constitutive or co-regulated with expression of genes targeting cellulose. The ability of *T. turnerae* to express these activities in the absence of induction by their specific target substrates is consistent with the observation that shipworm endosymbionts express a wide range of lignocellulolytic enzymes when growing within symbiont-containing cells (bacteriocytes) in the gill of the host where they have no direct contact with lignocellulose (O’Connor *et al*., 2014; Sabbadin *et al*., 2018).

In addition to proteomic predictions, we show that these MV preparations can hydrolyze CMC with specific activities comparable to or greater than that observed for a commercial purified cellulase enzyme cocktail (Figure 4 C). Although CMC is not cleaved by all cellulolytic enzymes, hydrolysis of CMC confirms that at least some of the detected cellulases retain their activity in purified MVs.

### TonB-dependent receptors in *T. turnerae* MVs

Although CAZymes constitute a substantial proportion of *T. turnerae* MV proteomes, the most abundant proteins observed in MV preparations were identified as TonB-dependent receptors (TBDRs). As previously reported in MVs of several Gram-negative bacteria (Veith *et al*., 2015; Zakharzhevskaya *et al*., 2017; Dhurve *et al*., 2022; Fadeev *et al*., 2023), TBDRs account for nearly half of the proteins detected in *T. turnerae* MVs. TBDRs are outer membrane-associated proteins that bind and mediate the energy-dependent movement of siderophores and various nutrients, including polysaccharides too large to be taken up via transmembrane diffusion across the outer membrane. Recently, it was shown that two of *T. turnerae’s* four *TonB* genes are essential for growth in iron-limiting conditions and for growth with cellulose as a sole carbon source, indicating that the *TonB* system (TBDR and *TonB*) is necessary for both iron acquisition and cellulose catabolism by *T. turnerae* (Naka and Haygood, 2023).

Interestingly, in a proteomic examination of the shipworm *Banka setacea*, a TBDR was the only major symbiont-derived protein observed in the cecum content that was not associated with the decomposition of lignocellulose (O’Connor *et al*., 2014). This finding suggests the presence of symbiont outer membranes in the cecum. However, the cecum of shipworms has been shown by microscopic (Betcher *et al*., 2012; Pesante *et al*., 2021), transcriptomic (Sabbadin *et al*., 2018), and proteomic (O’Connor *et al*., 2014) methods to be nearly devoid of intact bacterial cells. These observations argue against the transport of whole bacterial cells from the gill to the cecum, as suggested by (Pesante *et al*., 2021), but might suggest a role for MVs in transporting lignocellulose degrading enzymes from the gill to the gut in shipworms.

### Roles of MVs in symbiotic associations

Membrane vesicles are critical in bacterial interactions with many animal and plant hosts (Berleman and Auer, 2013; Lynch and Alegado, 2017). While most research has focused on pathogenic interactions (Gan *et al*., 2023), MVs may also contribute to beneficial host-symbiont associations. For example, MVs have been shown to mimic whole cells of *Vibrio fisheri* in the induction of specific aspects of light organ development in the Hawaiian bobtail squid *Euprymna scolope* (Aschtgen *et al*., 2016). Similarly, bacterial MVs significantly promoted the settlement of marine sponge larvae (Zhang *et al*., 2024). MVs are also produced by chemoautotrophic symbionts that occur in the trophosome of flatworms of the genus *Paracatenula,* where they play a critical role in providing the host with carbohydrates synthesized by the symbionts (Jäckle *et al*., 2019).

### Potential role of MVs in biomass conversion

In addition to their potential roles in shipworm symbiosis, cellulolytic MVs are of significant interest in converting plant biomass into renewable liquid fuels or fine chemicals (Thakur *et al*., 2023). A significant challenge in biomass conversion is understanding how cooperation among cellulases and associated enzymes can improve the design of efficient enzyme cocktails tailored to individual feedstocks (Østby *et al*., 2020). MVs produced by *T. turnerae* and other lignocellulolytic organisms may represent nature-based solutions that reveal specific combinations, concentrations, and spatial organization of cooperating enzymes that significantly enhance lignocellulose degradation in nature and so might inspire the design of engineered lignocellulose deconstruction systems.

## Author Contribution Statement

**Mark T. Gasser:** Conceptualization; investigation; formal analysis; methodology; writing – original draft preparation. **Annie Liu:** Investigation; methodology. **Marvin A. Altamia:** Investigation; methodology. **Bryan R. Brensinger:** Investigation; methodology. **Sarah L. Brewer:** Investigation; methodology. **Ron Flatau:** Investigation; methodology. **Eric R. Hancock:** Investigation; methodology. **Sarah P. Preheim:** Supervision; writing – review and editing. **Claire Marie Filone:** Funding acquisition; supervision; writing – review and editing. **Dan L. Distel:** Funding acquisition; Conceptualization; supervision; writing – original draft preparation; writing – review and editing.

## Acknowledgments

We thank the PennVet Extracellular Vesicle Core SCR_022444 (Philadelphia, Pennsylvania) for work and assistance with MV isolation and NTA analysis; the University of Maryland School of Medicine’s and School of Dentistry’s Electron Microscopy Core Imaging Facility – EMCIF (Baltimore, Maryland) for work and assistance with TEM imaging; and the NYU Langone’s Proteomics Laboratory SCR_017926 (New York, New York) for work and help with mass spectrometry analysis.

## Supplemental Information

Figure_S1_SuppInfo.docx

Dataset_S1_SuppInfo.xlsx

