## Supporting Information: Figure S1 for "Membrane vesicles can contribute to cellulose degradation by *Teredinibacter turnerae,* a cultivable intracellular endosymbiont of shipworms"

### Supplemental Figures

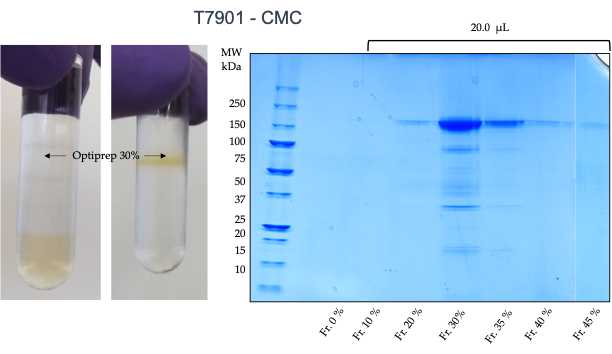

**B)**

**A)**

**After**

**Before**

Figure S1. Purification of T. turnerae crude OMVs by density gradient ultracentrifugation. (A) Representative image of density gradient banding before centrifugation and sample visualization at the interface of 30% fraction. (B) Representative SDS-PAGE analysis of OMVs collected from each density fraction.
